# Phylogenetic diversity of 200+ isolates of the ectomycorrhizal fungus *Cenococcum geophilum* associated with *Populus trichocarpa* soils in the Pacific Northwest, USA and comparison to globally distributed representatives

**DOI:** 10.1101/2020.03.24.005348

**Authors:** Jessica M. Vélez, Reese M. Morris, Rytas Vilgalys, Jessy Labbé, Christopher W. Schadt

## Abstract

The ectomycorrhizal fungal symbiont *Cenococcum geophilum* is of great interest as it is globally distributed, associates with many plant species, and has resistance to multiple environmental stressors. *C. geophilum* is only known from asexual states but is often considered a cryptic species complex, since extreme phylogenetic divergence is often observed within morphologically identical strains. Alternatively, *C. geophilum* may represent a highly diverse single species, which would suggest cryptic but frequent recombination. Here we describe a new isolate collection of 229 *C. geophilum* isolates from soils under *Populus trichocarpa* at 123 collection sites spanning a ~283 mile north-south transect in Washington and Oregon, USA (PNW). To further understanding of the phylogenetic relationships within *C. geophilum*, we performed maximum likelihood phylogenetic analyses to assess divergence within the PNW isolate collection, as well as a global *GAPDH* phylogenetic analysis of 789 isolates with publicly available data from the United States, Europe, Japan, and other countries. Phylogenetic analyses of the PNW isolates revealed three distinct phylogenetic groups, with 15 clades that strongly resolved at >80% bootstrap support based on a *GAPDH* phylogeny and one clade segregating strongly in two principle component analyses. The abundance and representation of PNW isolate clades varied greatly across the North-South range, including a monophyletic group of isolates that spanned nearly the entire gradient at ~250 miles. A direct comparison between the *GAPDH* and ITS phylogenies combined with additional analyses revealed stark incongruence between the ITS and *GAPDH* gene regions, implicating probable intra-species recombination between PNW isolates. In the global isolate collection phylogeny, 34 clades were strongly resolved at >80% bootstrap support, with some clades having intra- and intercontinental distributions. These data are highly suggestive of divergence within multiple cryptic species, however additional analyses such as higher resolution genotype-by-sequencing approaches are needed to distinguish potential species boundaries and recombination patterns.

## Introduction

Plant-fungal relationships are often difficult to disentangle, as a single plant species may be associated with hundreds of fungal species and each of these associations can have varying influences on host plant survival and growth that co-vary with the environmental and physical conditions of soil (Bonito et al., 2014; Grayston et al., 1998; Kennedy et al., 2011; Raynaud et al., 2008). The complexity of these plant-fungal relationships may be at least partially illuminated through targeted understanding of the interactions of exemplar fungal species. Such model species can serve as representatives for exploring plant-fungal interactions and characteristics across various conditions. The genus *Populus* is an excellent plant model for such studies because *Populus trichocarpa* has a fully sequenced genome (Torr et al., 2006; Tuskan et al., 2004) and hosts a diverse community of microbes including bacteria, archaea and fungi (Bonito et al., 2014; Cregger et al., 2018; Hacquard and Schadt, 2015) which are capable of accessing, metabolizing, producing, and/or immobilizing compounds which the plant cannot (Berendsen et al., 2012). With over 30 species of deciduous, fast-growing softwoods with distinct sexes and natural hybridization (Jansson et al., 2010), *Populus* also has great importance in agroforestry industries as pulp and paper, and potential bioenergy feedstocks (Porth and El-Kassaby, 2015; Sannigrahi et al., 2012).

Similar to *Populus* spp., a correspondingly robust model ectomycorrhizal (ECM) fungal group for paired studies with *Populus* should be widespread, interact with many plant hosts, and be easily culturable in a laboratory setting. The ECM fungal species *Cenococcum geophilum* is a ubiquitously distributed fungus which is positively associated with plant health, growth, and increased soil contaminant resistance (Meena et al., 2017; Zong et al., 2015), and as such has the potential to serve well as a model organism for genetic, physiological, and ecological studies. *Cenococcum geophilum* is known to associate with both angiosperm and gymnosperm species across 40 plant genera, representing over 200 host species (Douhan et al., 2007b), and is generally tolerant across salinity gradients (Matsuda et al., 2017), under water stress conditions (Fernandez and Koide, 2013), and in extreme inorganic soil contamination conditions (Fernandez and Koide, 2013; Krpata et al., 2008; Matsuda et al., 2017). This wide-ranging resistance to stress conditions in the soil environment is often associated with a high melanin content (Cordero and Casadevall, 2017; Fernandez and Koide, 2013; Matsuda et al., 2017; Toledo et al., 2017) which may also contribute to longevity and resistance to decomposition in soils (Fernandez et al., 2013). *Cenococcum geophilum* is readily culturable in a laboratory setting and capable of growing on multiple standard solid media, including both defined media such as Modified Melin-Norkrans (MMN) and complex media such as potato dextrose agar (PDA). Additionally, laboratory methods for targeted and relatively rapid isolation from soils via its abundant and phenotypically characteristic sclerotia (Obase et al., 2014; Trappe, 1969) allow for efficient isolation of new strains directly from soil samples. All of these characteristics make *C. geophilum* an ideal species for population-level genomic studies. However most existing regional-level isolate collections focus primarily on coniferous species (de Freitas Pereira et al., 2018; Matsuda et al., 2015; Obase et al., 2016) rather than angiosperms such as *Populus*.

The first sequenced genome of *C. geophilum* strain 1.58 (isolated from Switzerland) was published along with the Joint Genome Institute (JGI) in 2016 (Peter et al., 2016). This genome is among the largest in the fungal kingdom, with a mapped size of 178 Mbp and a total estimated size of up to 203 Mbp (Peter et al., 2016; Talhinhas et al., 2017). *Cenococcum geophilum* has no documented means of sexual spore production (Douhan et al., 2007b, 2007c; Jany et al., 2002), and is considered asexual as a species despite high levels of genetic and physiological diversity. Due to high levels of diversity that are reported among *C. geophilum* isolates, even within isolates from beneath a single tree (Bahram et al., 2011), *C. geophilum* has been suggested to represent a complex of indistinguishable cryptic species (Douhan et al., 2007b; Matsuda et al., 2015; Obase et al., 2017; Peter et al., 2016), with many studies finding significant variation in cultured isolate characteristics and physiology (Douhan et al., 2007c). However, patterns supporting recombination have also been observed by previous studies, suggesting a cryptic sexual state or other mechanisms for intrapopulation recombination (Bourne et al., 2014; Douhan et al., 2007c; Jany et al., 2002; LoBuglio and Taylor, 2002; Peter et al., 2016). Together these studies have led many to suggest that *Cenococcum* represents an unknown number of cryptically separate species on the genetic level (Douhan et al., 2007c, 2007b; Douhan and Rizzo, 2005). Further supporting this suggestion, a 2016 multigene phylogeny of a previously characterized *C. geophilum* isolate collection (Obase et al., 2014) revealed a divergent clade described as *Pseudocenococcum floridanum* (Obase et al., 2016). This discovery highlights the need to further explore the phylogenetic diversity among diverse regional isolates of *C. geophilum* in order to better characterize this species and determine several factors, including 1) if *C. geophilum* is a single highly outcrossed species, or a variety of cryptic species; 2) if a variety of species, how diverse the populations are and what are the patterns of speciation; and 3) if there exists intra- or interspecies patterns of recombination.

*Cenococcum geophilum* thus appears to have a myriad of potential benefits as a model ECM and rhizosphere-associated species. However, this fungus has been difficult to study phylogenetically, and further work is needed to delineate its phylogenetic and functional relationships within what may be a highly heterogenous species complex. Our laboratory set a long-term goal to build a genetically diverse collection of *C. geophilum* isolates from across the *Populus trichocarpa* range, mirroring the host GWAS population studies which have proven valuable for understanding the biology of these host trees (Bdeir et al., 2019; Evans et al., 2014; Franco-Coronado et al., 2018; Mckown et al., 2014; McKown et al., 2017; Muchero et al., 2011; Tuskan et al., 2018; Zhang et al., 2018). Towards this goal we have isolated 229 new *C. geophilum* strains from beneath *P. trichocarpa* stands over a range of approximately 283 miles of the host range in the United States Pacific Northwest (PNW) states of Oregon and Washington and confirmed their identity using sequencing of the internal transcribed spacer (ITS) region and the glyceraldehyde-3-phosphate dehydrogenase (*GAPDH*) gene. Maximum likelihood phylogenies of the *GAPDH* gene and ITS region of our PNW collection were compared directly in order to identify potential patterns of intra-species recombination. Furthermore, the *GAPDH* genes of our PNW collection were additionally compared to over 500 *C. geophilum* isolates with available data from published isolates gathered from other locations and studies primarily in the United States, Europe, Japan, but also other sites where data were publicly available (de Freitas Pereira et al., 2018; Douhan et al., 2007b; Douhan and Rizzo, 2005; Matsuda et al., 2015; Obase et al., 2016, 2014), as well as 16 additional European isolates recently sequenced by JGI and provided by Drs. Francis Martin and Martina Peter (Freitas Pereira et al., 2018).

## Materials and Methods

### Site selection and soil sampling

Primary sampling was carried out over a six-day period in late July of 2016. At each site, three one-gallon soil samples were collected directly under *P. trichocarpa* with at least 50 yards distance in between each collection. Typically, at least three sampling sites were selected within each watershed from the Willamette River in Central Oregon to the Nooksack river near the Canadian border along a north-south gradient (i.e. the Interstate 5 corridor). Several watersheds and sites corresponded to those sampled in Evans et. al. (2014) where possible to allow for testing of associations with *P. trichocarpa* GWAS populations in future studies. Soil samples were collected to approximately a 20 cm depth. A trowel was used to fill one-gallon Ziploc freezer bags which were kept on ice and refrigerated at 4°C until analysis. Site soil temperatures were recorded using an electronic thermometer (OMEGA model RDXL4SD). Upon return to the lab, two 15 mL tubes of the 105 samples in 2016 were subsampled for soil moisture, carbon (C), nitrogen (N), and soil elemental characterization. Additionally, several isolates derived from smaller exploratory soil samples from within the southern end of this geographic range (sampled by Drs. R. Vilgalys and C. Schadt) for methods development during the prior year were also included in this study, for a total of 123 soil samples.

### Sclerotia separation

Soil samples were prepared using a procedure described by Obase et al. (2014) with modifications. Samples were manually sieved and rinsed with distilled water to retain particles between 2mm and 500 μM, and the resulting slurry allowed to soak in distilled water for 10-30 seconds. Floating debris could be decanted off the top. Portions of the slurry were placed into gridded square Petri dishes (VWR 60872-310) which were partially filled with water and a dissection scope was used to remove sclerotia using tweezers. Sclerotia were submerged in undiluted Clorox bleach for 40 minutes using a VWR 40 μm nylon cell strainer (Sigma-Aldrich Z742102), then rinsed 3 times with sterile distilled water. Sclerotia which did not turn white after bleach treatment were plated onto Modified Melin-Norkrans (MMN) media (Marx, 1969) composed of 3 g l^−1^ malt extract, 1.25 g l^−1^ glucose, 0.25 g l^−1^ (NH_4_)_2_HPO_4_, 0.5 g l^−1^ KH_2_PO_4_, 0.15 g l^−1^ MgSO_4_-7H_2_O, 0.05 g l^−1^ CaCl_2_, 1 mL^−1^ FeCl_3_ of 1% aqueous solution, and 10 g l^−1^ agar, adjusted to 7.0 pH using 1N NaOH. After autoclaving media were allowed to cool to 60°C, then 1 g l^−1^ thiamine was added along with the antibiotics Ampicillin and Streptomycin at 100 ppm each. Plates with sclerotia were stored in the dark at 20°C.

### DNA extraction

Isolates with dark black growth were considered viable and allowed to grow until ~5 mm diameter. Colonies were transferred onto a cellulose grid filter (GN Metricel 28148-813) on MMN plates using the previous recipe except adding 7 g l^−1^ dextrose and omitting antibiotics and allowed to grow for 1-3 months for DNA extraction. The Extract-N-Amp kit (Sigma-Aldrich XNAP2-1KT) was used as per manufacturer instructions to extract genomic DNA, with the modification to use 20 μL of the Extraction and Dilution solutions (Kluber et al., 2012). DNA samples were stored at −20°C until use.

### PCR amplification

Ribosomal DNA (rRNA) was amplified using fungal-specific ITS primers ITS1 and ITS4 (Bokulich and Mills, 2013), and the *GAPDH* gene was amplified using the gpd1 and gpd2 primers (Berbee et al., 1999) using the Promega GoTaq © Master Mix kit to amplify DNA. The thermocycling conditions consisted of an initial hold of 94°C for 5 min, followed by 35 cycles of 94°C (30 s), 55°C (30 s), and 72°C (2 min), with a final elongation of 72°C for 10 min. Amplified PCR products were analyzed on a 1% agarose gel using TAE buffer to confirm band size prior to cleanup. Samples were then cleaned using the Affymetrix USB ExoSAP-IT © kit and sequenced on an ABI3730 Genetic Analyzer at the University of Tennessee at Knoxville (UTK), or at Eurofins Genomics (Louisville, Kentucky, USA). Sequences generated were analyzed against the NCBI database using the BLAST feature in Geneious version 10.2.3 (https://www.geneious.com, Kearse et al., 2012) to verify fungal identity as *C. geophilum*. Confirmed *C. geophilum* isolates (marked “CG” with number) were stored on MMN plates at 20°C and re-plated quarterly to ensure continued viability.

### Determination of native soil properties

For the 105 soil samples collected in July of 2016, soil temperature, concentrations of carbon, nitrogen, elemental metals, and soil water content were analyzed. A C/N analysis was conducted on approximately 18g of each soil sample. The samples were oven-dried at 70°C and ground to a fine powder. Approximately 0.2 g of ground sample were analyzed for carbon and nitrogen on a LECO TruSpec elemental analyzer (LECO Corporation, St. Joseph, MI). Duplicate samples and a standard of known carbon and nitrogen concentration (Soil lot 1010, LECO Corporation, carbon = 2.77 % ± 0.06 % SD, nitrogen = .233 % ± 0.013 % SD) were used to ensure the accuracy and precision of the data.

Soil elemental metal concentrations were determined using the Bruker Tracer III-SD XRF device. Approximately 1g of dried, homogenized soil was placed into Chemplex 1500 series sample cups with Chemplex 1600 series vented caps and 6 uM Chemplex Mylar® Thin-Film. Cups were placed against the XRF examination window and scanned for 60 seconds at 40 kV with a vacuum and no filter. Elemental spectra were collected using the Bruker S1PXRF S1 MODE v. 3.8.30 software and analyzed using the Bruker Spectra ARTAX v. 7.4.0.0 software.

Correlation coefficients relating the total sclerotia isolated, total *Cenococcum* isolates per site, soil temperature, percent moisture content, C and N weight percent, and total counts per minute (cpm) of a range of elements were then calculated using the R v.3.4.1 statistical analysis software (2017) and corrplot package v. 0.84 (https://cran.r-project.org/web/packages/corrplot/index.html) in order to determine whether relationships existed between soil quality and content and *C. geophilum* abundance and isolation success.

### Generation of phylogenies

The ITS 5.8s subunit and *GAPDH* gene sequences of 228 PNW *C. geophilum* isolates were successfully amplified and sequenced, trimmed and aligned to the published genome of the *C. geophilum* strain 1.58 using Geneious. The ITS and *GAPDH* phylogenies were concatenated using Geneious, and isolates lacking either ITS or *GAPDH* sequence data were excluded from the multigene concatenated analysis. For comparison with globally distributed isolates, ITS and *GAPDH* sequence data of 543 *C. geophilum* strains were obtained from GenBank including strains from Japan, Europe, the United States, and 16 additional European isolates recently sequenced by JGI were provided by Drs. Francis Martin and Martina Peter (personal communication, Freitas Pereira et al., 2018). These isolates were aligned with the Pacific Northwest (PNW) isolate collection using ITS and *GAPDH* separately, as well as a multigene concatenation of ITS and *GAPDH*. Isolates lacking either ITS or *GAPDH* sequence data were excluded from the concatenated ML analysis of the global isolate collection for a total of 499 global collection isolates. All phylogenies were rooted using the outgroups *Glonium stellatum, Hysterium pulicare*, and *Pseudocenococcum floridanum* isolate BA4b018 (Obase et al., 2016), which were downloaded from either GenBank (https://www.ncbi.nlm.nih.gov/genbank/) or MycoCosm (Grigoriev et al., 2014; Nordberg et al., 2014) (https://genome.jgi.doe.gov/programs/fungi/index.jsf).

### Comparison of GAPDH, ITS, and GAPDH + ITS phylogenies

The best models for maximum likelihood analyses were determined for each individual gene alignment and combined gene region alignments using the Find Best DNA/Protein Models feature in MEGA X (Kumar et al., 2018) to determine the Akaike Information Criterion (AICc) value (Akaike, 1973) and Bayesian Information Criterion (BIC) (Schwarz, 1978), with the best model indicated by the lowest AICc and BIC values (Table 1). A maximum likelihood (ML) phylogenetic analysis was produced for the PNW isolate collection ITS, *GAPDH* and concatenated phylogenies using the MEGA X software with the determined best model settings using 1000 bootstrap replications. The produced ITS and *GAPDH* ML analyses of the PNW isolate collection were directly compared to both the concatenated phylogeny separately, and to each other, using the TreeGraph2 software (Stöver and Müller, 2010) and the R v.3.4.1 statistical analysis software. These analyses indicated disagreement between the two gene datasets and thus only *GAPDH* was used for further analyses as it contained the most phylogenetically informative sites.

**Table 1.** Model with best fit analysis, Bayesian Information Criterion, and Akaike Information Criterion (AICc) for each alignment per an analysis using MEGA X software. Lower BIC and AICc values indicate the best model fit for use in analyses.

| Alignment | Best Model | AICc | BIC |
| --- | --- | --- | --- |
| PNW ITS | K2 + G | 2873.433 | 7082.611 |
| PNW *GAPDH* | K2 + G | 5468.328 | 9860.315 |
| PNW ITS + *GAPDH* | K2 + G | 8322.607 | 12970.723 |
| GP ITS | K2 + G | 7258.62 | 21156.093 |
| GP *GAPDH* | K2 + G | 12807.63 | 29779.549 |
| GP ITS + *GAPDH* | K2 + G + I | 15623 | 26550.792 |

### Phylogeographic variation within PNW isolates

A correlation plot was created using R statistical software packages Hmisc v. 4.2.0 and corrplot v. 0.84 to determine whether resolved clades correlated with the latitude of the strain isolation site, and a multiple correspondence analysis (MCA) was conducted to determine clade-specific correlations to latitude. Additionally, a principle components analysis (PCoA) was conducted using a distance matrix generated from the PNW *GAPDH* phylogeny with and without species outgroups included in order to determine potential speciation patterns within the PNW isolate collection. The PNW ML analysis was mapped by isolate site latitude to the PNW region using the GenGIS 2.5.3 software (Parks et al., 2013) in order to visualize spatial diversity within the PNW isolates and designated clades.

### Global GAPDH phylogenetic analyses

In order to directly compare analyses of the PNW and global isolate collections, the global *GAPDH* phylogeny was constructed using a maximum likelihood (ML) analysis in the MEGA X software (Kumar et al., 2018) with 1000 bootstrap replications and the previously determined best fit settings (Table 1). Phylogenetic trees were visualized indicating either bootstrap support values or branch lengths using FigTree v. 1.4.4 (http://tree.bio.ed.ac.uk/software/figtree/). Clades were designated as strongly grouped isolates with bootstrap support values of >80%.

In addition to avoiding conflicts between *GAPDH* and ITS, usage of the *GAPDH* rather than multigene concatenated phylogeny allowed for a more comprehensive global isolate collection analysis, as the concatenated global isolate phylogeny excluded >300 isolates which lacked both sequences while the *GAPDH* global isolate phylogeny included a total of 789 isolates. As a result, only the *GAPDH* dataset was used for global collection analyses.

### Recombination patterns within GAPDH and ITS data

The ITS and *GAPDH* RAxML phylogenies were visually compared and analyzed using R package phytools v. 0.6.99. A PCoA was completed for the ITS, *GAPDH*, and concatenated phylogenies and alignments using distance matrices generated using the ape v. 5.3 and seqinr v. 3.6-1 packages in the R statistical software in order to compare patterns of recombination within the PNW isolate collection based on either or both gene regions. Additionally, HKY85 distance matrices were generated using Phylogenetic Analysis Using Parsimony (PAUP) v. 4 (Swofford, 2003) and graphed against each other in the R statistical software using the ggplot2 package (Wickam, 2016). The inter-partition length difference (ILD) test was used to assess phylogenetic congruence between ITS and *GAPDH* data sets for the PNW isolates using the PTP-ILD option in PAUP* Version 4 (Swofford, 2003), with 1000 permutations and default settings.

## Results

### A diverse culture collection of PNW Cenococcum isolates associated with Populus

A total of 229 PNW isolates were obtained from 56 out of 123 soil samples (105 primary soil samples from 2016 + 18 preliminary samples from 2015 – Supplemental data, Table 1), accounting for a 46% overall *C. geophilum* isolation success rate, as some soil samples had few or no sclerotia. The 105 PNW primary soil samples for which associated soils data were also generated, the total *C. geophilum* isolation success rate did not positively correlate to any measured soil condition or quality (Supplemental Fig 1). Isolation success also did not strongly correlate to the total number of sclerotia recovered (r = 0.38, p >0.05). Between the soil values however there were expected relationships. For example, the strongest correlation was between the soil C and N weight percentages (r = 0.99, p = <0.05). Strong correlations also existed between the percent moisture content and C content (r = 0.88, p = <0.05) or N content (r = 0.86, p = <0.05), C content and zinc counts per minute (cpm) (r = 0.80, p = <0.05), and N content and zinc cpm (r = 0.81, p = <0.05) (Supplemental Fig 1).

A total of 438 *GAPDH* positions were represented in the alignment for the *GAPDH* ML analyses of the PNW and global isolate collections. In the PNW isolates, the phylogeny backbone strongly resolved the PNW collection at 97.1%. A total of 155 isolates grouped into 15 clades where two or more strains were resolved at >80% bootstrap values (Fig 1, Supplemental data, Table 2). Of the 15 resolved clades, two contain nested clades of two or more isolates which resolved at >80% bootstrap support (Supplemental data, Table 2). Nonetheless, 74 of our 229 PNW *Populus* isolates were not strongly resolved by these analyses.

**Fig 1:**
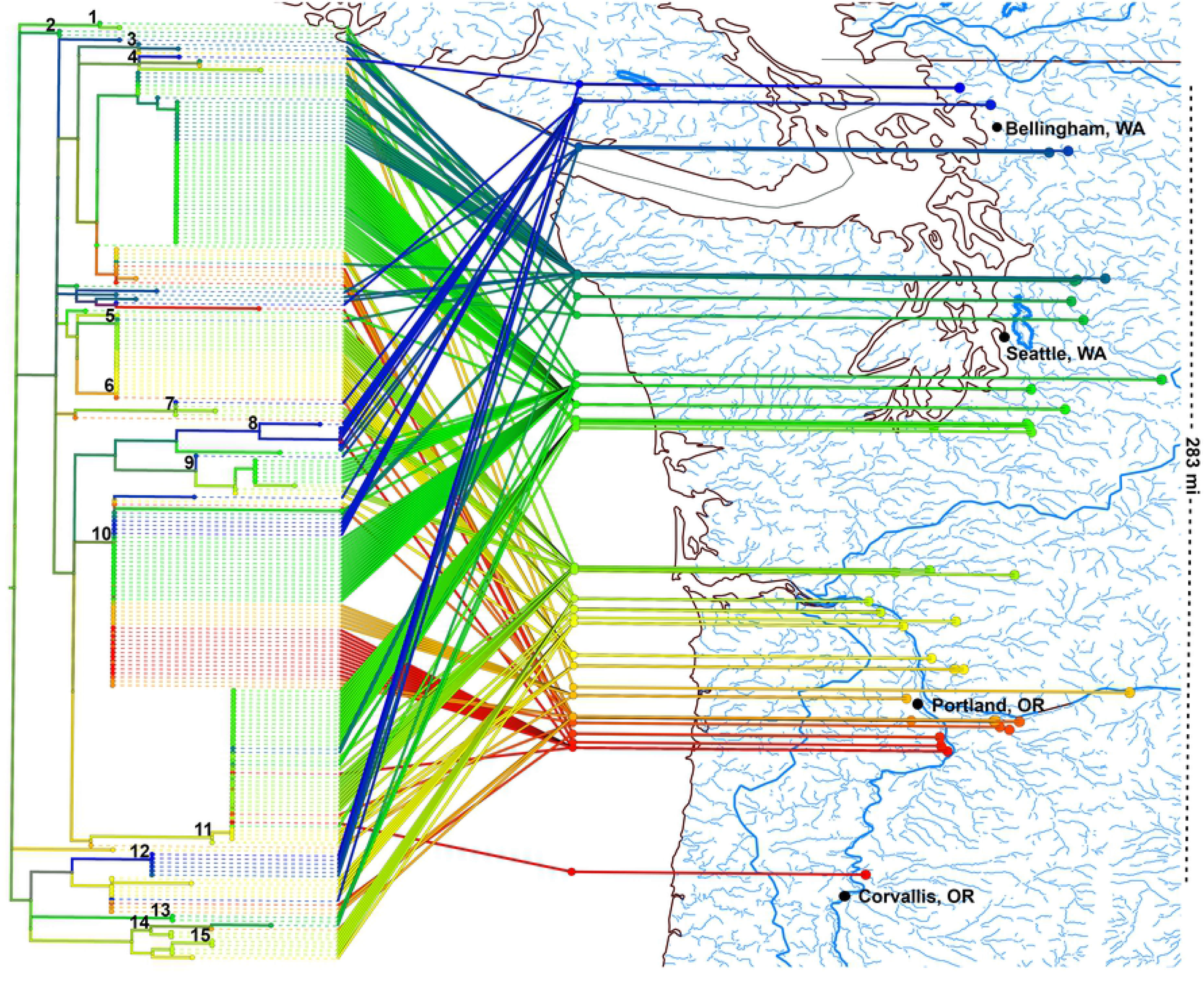
The Pacific Northwest (PNW) isolate *GAPDH* RAxML phylogeny mapped by latitude to the PNW region. Clades with >80% bootstrap support values are indicated above the associated clade. A total of 15 clades encompassing 155 isolates were identified in the PNW isolate collection, with 74 isolates remaining unresolved. Isolates from smaller clades tended to group by region of origin with a few notable exceptions present over large north-south distances.

### Phylogeographic variation within PNW isolates

Phylogenetic distance matrices analyzed via PCoA revealed that the PNW isolates group separately as three distinct groups. When using a distance matrix based on the RAxML phylogeny, the PNW 11 clade segregates as a distinct phylogenetic group (Fig 2A), and when using an alignment-generated distance matrix, clades 10 and 11 both segregate as distinct phylogenetic groups (Fig 2B).

**Fig 2:**
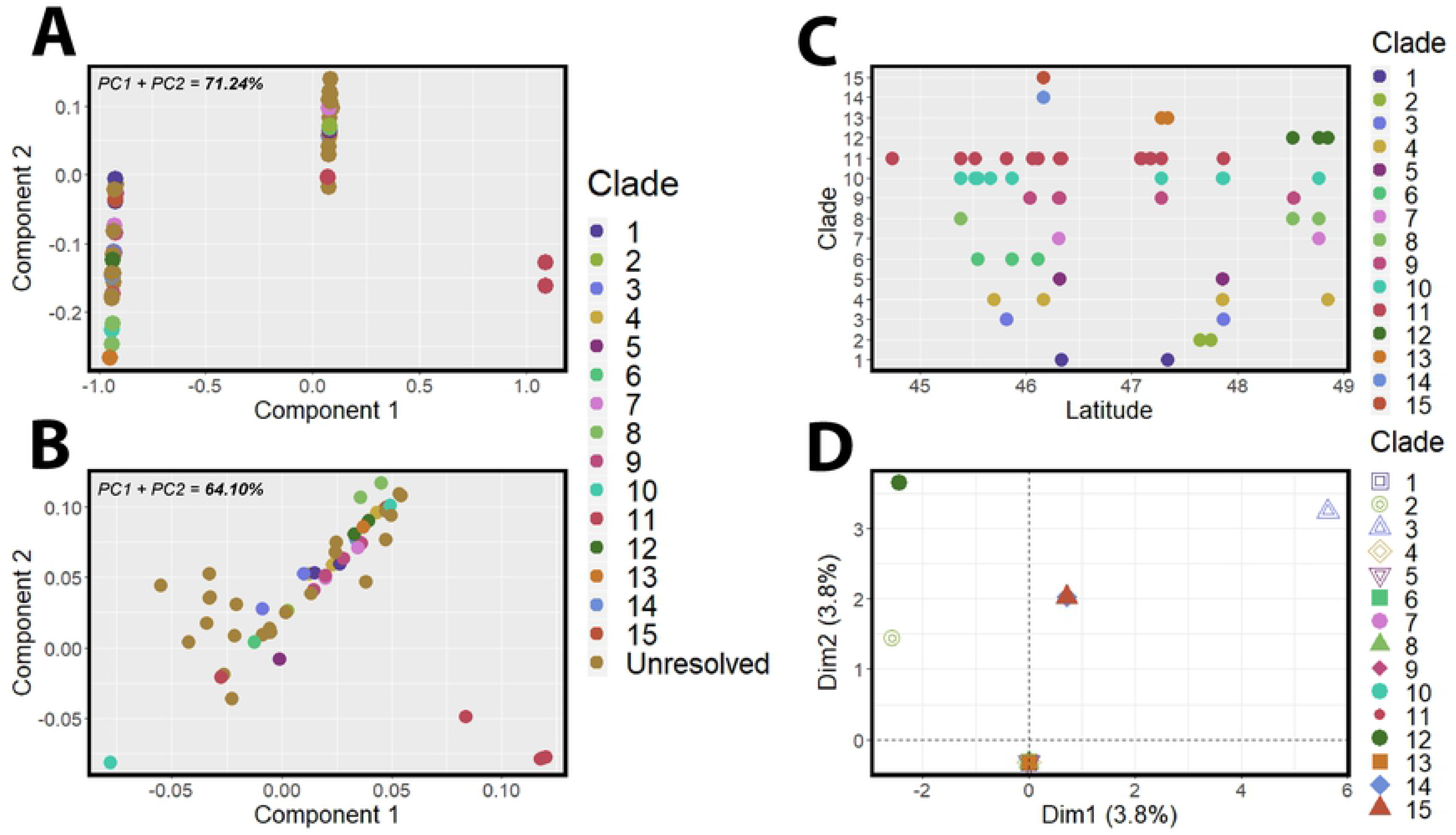
Principle components analysis (PCoA) of the PNW *GAPDH* RAxML phylogeny revealed three distinct isolate clusters, with clade 11 segregating as a separate cluster from the remaining PNW collection (A). A PCoA of the PNW *GAPDH* alignment also revealed three distinct clusters, with clades 10 and 11 segregating from the remaining PNW collection (B). A scatterplot of clade versus latitude did not reveal distinct patterns within the larger groups of the PNW collection (C), and a multiple components analysis showed some differentiation with clades 3, 12, 14, and 15 from the remaining PNW isolate collection, but revealed weak associations overall between latitude, isolate clade, and phylogenetic differentiation (D).

The *GAPDH* phylogenetic analysis mapped to the PNW soil sampling sites by latitude revealed that isolates did not appear to strongly correlate to original isolation site latitudes (Fig 1, Supplemental data, Table 2). High geographic latitude variation is seen within clade 11, which includes 38 strains isolated >60 miles apart (Fig 1, Supplemental data, Table 2). Clades three, four, five, seven, and nine include strains isolated >100 miles apart (Fig 1, Supplemental data, Table 2). The nested clade 8.1 has the overall largest spatial range, with a single Willamette river isolate WI7_83.9 grouping with several isolates from sites >200 miles to the north (Fig 1, Supplemental data, Table 2) on the Nooksack River near the Canadian border. Clade ten is the largest clade with a total of 45 isolates (Fig 1, Supplemental data, Table 2), representing the greatest spatial diversity between individual sites within a single clade. The direct mapping to the origin of isolation latitude revealed no clear patterns of segregation, particularly in the largest clades. Further supporting this lack of association with latitude, an MCA found that the combination of clade and latitude were weakly associated with phylogenetic variation within the PNW isolate collection, with the combination of these two factors accounting for only 7.6% of the phylogenetic variation found (Fig 2D). A deeper analysis did reveal that clades three, 12, 14, and 15 are additionally not strongly grouped with the majority of the PNW isolate collection (Fig 2D).

### Recombination analyses in the PNW isolate collection

While both the ITS and *GAPDH* phylogenies of the PNW isolates had a strongly resolved backbone (>80% support), the PNW *GAPDH* phylogeny strongly resolved 15 clades within the PNW isolate collection while the PNW ITS phylogeny strongly resolved only 3 clades and showed numerous apparent conflicts (Fig 3A). When pairwise HKY85 distances of both genes were plotted against each other, a very low but still significant level of correlation was observed between the ITS and *GAPDH* gene regions (R2=0.185, p<.001, Fig 3B). To further validate this apparent incongruence between the two genes, a total of 102 informative sites of the concatenated PNW alignment were analyzed using the ILD in PAUP. The ILD test confirmed that the ITS and *GAPDH* data sets are incongruent (Fig 3C), as the summed lengths of the two parsimony trees made from the observed data set were significantly shorter than the distribution of combined tree lengths calculated for 1000 randomizations of the data set (p = 0.001).

**Fig 3:**
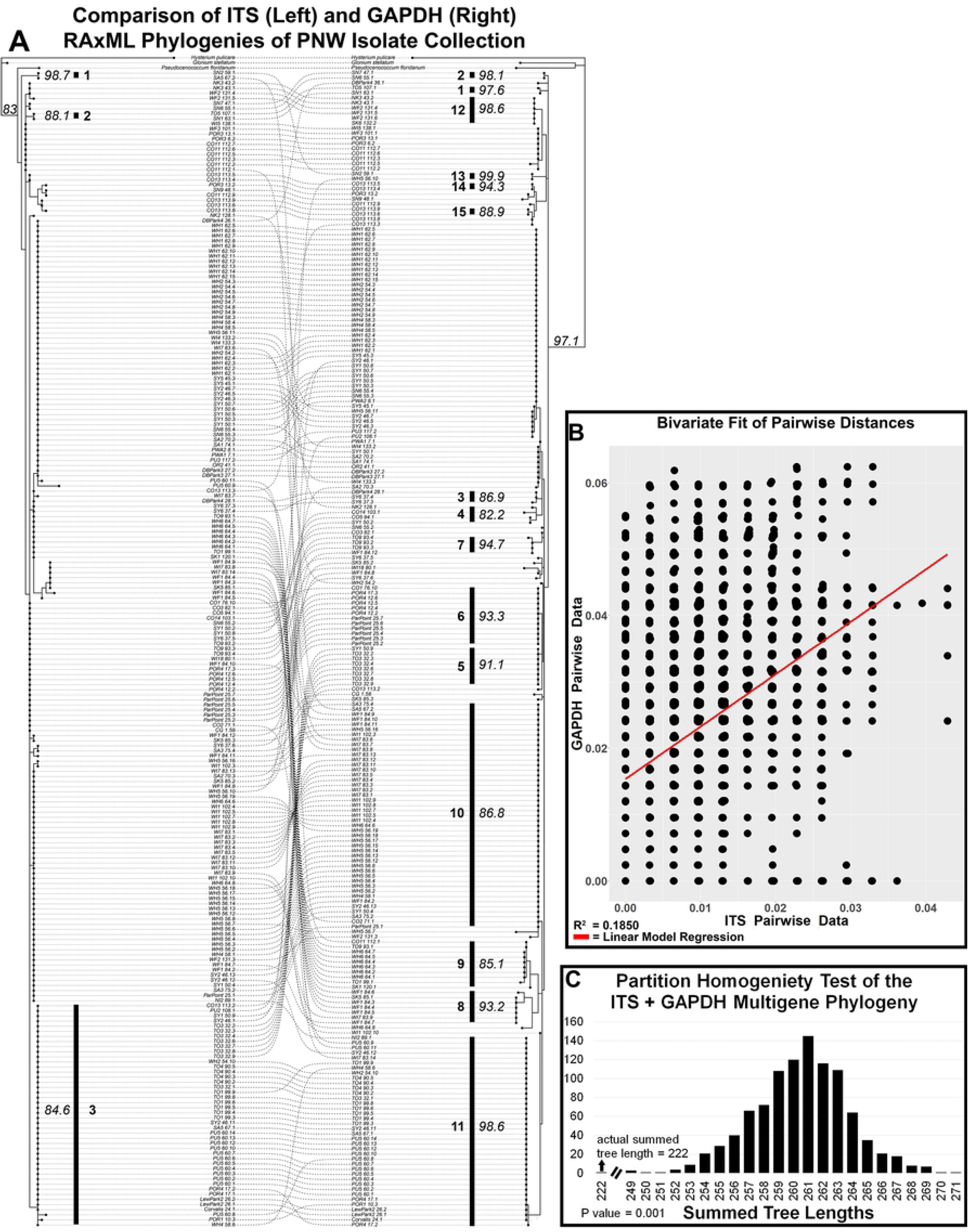
Phylogenetic incongruence between the ITS (left) and *GAPDH* (right) RAxML phylogenies (A). A scatterplot of pairwise HKY85 distances for ITS vs *GAPDH* datasets shows low correlation between the ITS and *GAPDH* gene regions (B). The parsimony-based inter-partition length difference (PTP-ILD)test indicated that the ITS and *GAPDH* data sets are incongruent due to no significant difference between the observed data set parsimony trees and the distribution of the randomized tree length calculations (p = 0.001) (C).

### Phylogeographic variation within the global C. geophilum isolate collection

Phylogenetic analyses of the PNW isolate collection together with the larger global isolate collection revealed 34 clades of two or more strains resolved at >80% bootstrap support (Fig 4, Supplemental data, Table 3). The major PNW clades 1, 2, 5, 6, 7, 11, 12, 13, 14, and 15 grouped identically in the global *GAPDH* analysis (clades V1, V2, V5, V6, V7, V11, V12, V13, V14, and V15 respectively – Fig 4, Supplemental data, Table 3), and nested clade 4.1 also persisted as global isolate collection clade V4.1 with a bootstrap value of 96.5% (Fig 4, Supplemental data, Table 3). The PNW clade 8 persisted as clade V8 and additionally included isolates CAA022, CAA006, CLW001 and CLW033 from the Florida/Georgia region (Obase et al., 2016) (Fig 4, Supplemental data, Table 3).

**Fig 4:**
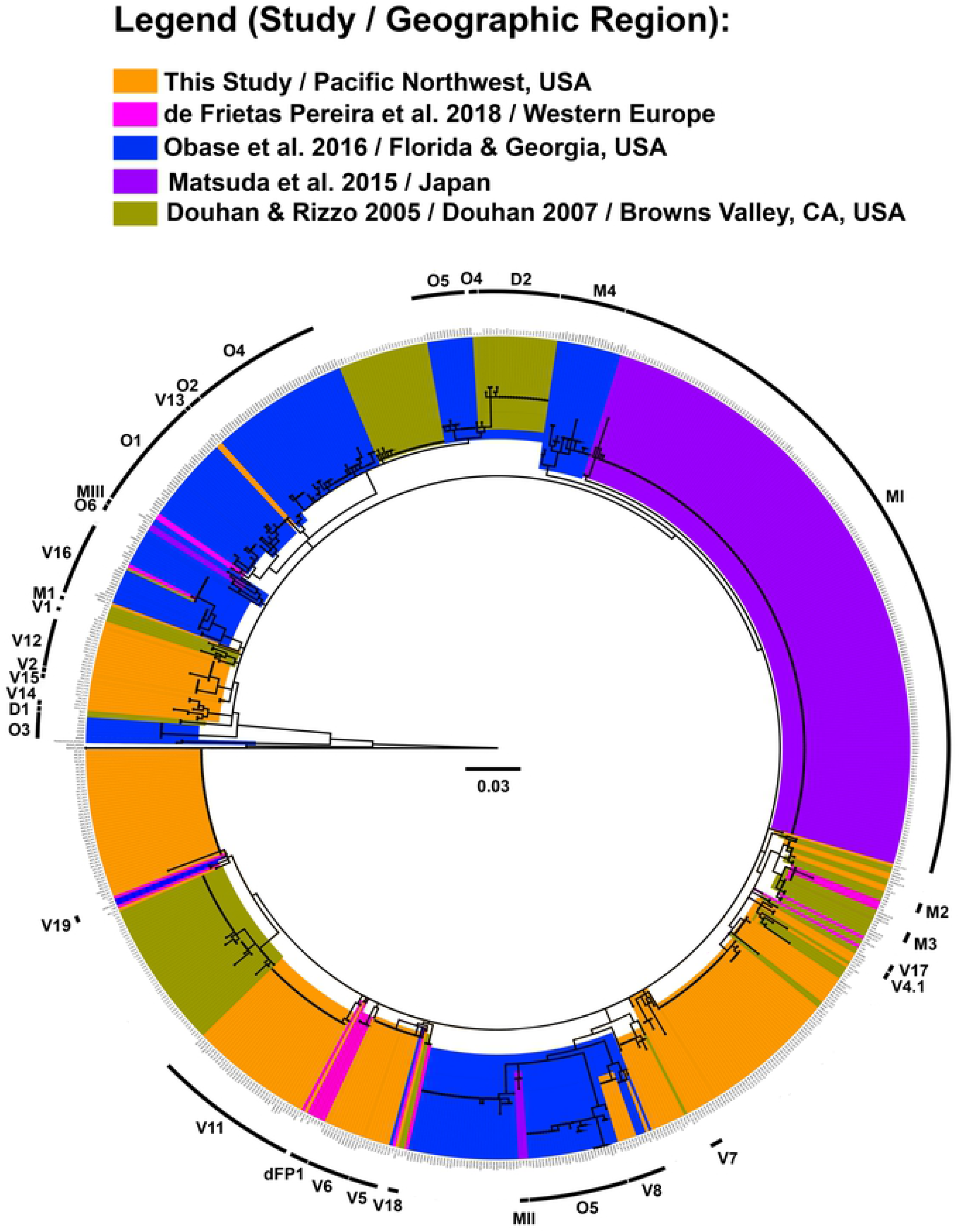
Global isolate collection *GAPDH* RAxML phylogenetic tree. Clades with strong bootstrap support (>80%) are labeled on the outer ring. Strongly supported clades implicated in this study are highlighted in orange and designated with V. Clades V16-V19 represent newly designated clades within the global *C. geophilum* isolate collection. The PNW isolates are highlighted in orange. Isolates are highlighted per the most recent published study of origin as follows: D (Douhan et al., 2007b; Douhan and Rizzo, 2005) highlighted in gold; dFP (de Freitas Pereira et al., 2018) highlighted in pink; M (Matsuda et al., 2015) highlighted in purple; and O (Obase et al., 2016) highlighted in blue.

## Discussion

### Cenococcum phylogeographic diversity in the Pacific Northwest

This study is the first comprehensive look at the genetic relationships within a regional population of *Cenococcum geophilum* isolates associated with a single host tree *Populus trichocarpa*. Previous *C. geophilum* genetic studies have primarily focused on gymnosperm species such as pines and Douglas fir (Matsuda et al., 2015; Obase et al., 2016), with two studies incorporating isolates collected under an angiosperm (oak) as well as other local gymnosperm host species (Douhan et al., 2007a; Douhan and Rizzo, 2005) and a second incorporating isolates collected under *Fagus sylvatica* (de Freitas Pereira et al., 2018) (Table 2). Interestingly, the genetic diversity encountered within this isolate collection exceeds the diversity observed in gymnosperm-associated populations by over twofold (Table 2) (Matsuda et al., 2015; Obase et al., 2016) implying that *P. trichocarpa* may exert fundamentally different selective pressures on the ECM *C. geophilum* than gymnosperm hosts.

**Table 2:** Previously resolved clades per study, number of isolates, geographic region, and host plant. The Pacific Northwest (PNW) isolate collection resolved 15 clades which appear to be uniquely associated with *Populus trichocarpa* within its host range in the PNW.

| Study | Geographic Region | Total No. Isolates | Host Plant | Gene(s) Used for Phylogenetic Analyses | No. clades resolved at >80% bootstrap support |
| --- | --- | --- | --- | --- | --- |
| Douhan & Rizzo 2005 | Browns Valley, California<br>Non-California (location not included) | 103<br>7 | Oak<br>Not indicated | *GAPDH* | 3 |
| Douhan & Huryn 2007 | Browns Valley, California<br>Pacific Northwest, USA<br>Alabama, USA<br>Alaska, USA<br>Europe | 74 | Mixed host species | *GAPDH* | 9<sup>c</sup> |
| Matsuda et al. 2015 | Japan | 225 | *Pinus thunbergii* | *GAPDH* | 3 |
| Obase et al. 2016 | Florida and Georgia, USA<br>Japan<br>Europe<br>North America | 242<sup>a</sup> | *Pinus elliotii*<br>*Pinus taeda* | ITS<sup>b</sup><br>*GAPDH*<sup>b</sup> | 7<sup>d</sup> |
| de Freitas et al. 2018 | Europe | 16 | *Picea abies*<br>*Pinus sylvestris*<br>*Fagus sylvatica*<br>Mixed forest | ITS<sup>b</sup><br>*GAPDH*<sup>b</sup> | 3<sup>e</sup> |
| PNW Collection | Pacific Northwest, USA | 231 | *Populus trichocarpa* | *GAPDH* | 15 |
| Global Collection | Pacific Northwest, USA<br>California, USA<br>Florida and Georgia, USA<br>Japan<br>Europe<br>North America | 790 | Mixed host species | *GAPDH* | 34 |

a: Representative isolates of 768 total
b: Concatenated
c: "...a high level of bootstrap support was not found for the majority of the backbone of the phylogeny and thus phylogeographic inference could not be made." (Douhan & Huryn, 2007)
d: Clade 7 delimited as novel species *P. floridanum*
e: Clades previously indicated in Obase et al. 2016

Based on a *GAPDH* sequence ML analysis, a total of 15 *C. geophilum* clades were resolved across a 283 mile transect of our sites selected along rivers in the PNW. Smaller clades tended to group latitudinally by site and watershed of origin, although notable exceptions exist, with two groups containing numerous nearly identical isolates despite distances of >100 miles between their sites of origin. In the PNW isolate collection, 74 isolates were outside of any other strongly supported clades (Fig 1, Supplemental Data, Table 2), but the majority grouped strongly, indicating potentially cryptic species represented by the strongly supported clades. Clade 10 and particularly clade 11, which consistently clusters separately from the remaining PNW isolate collection, may represent possible incipient species.

While geographic patterns are clearly evident within the phylogenetic analysis of our PNW isolates, interestingly the soil variables examined seemed to have no correlation with either the number of *C. geophilum* sclerotia recovered or the number of isolates obtained from our samples or their phylogenetic relationships (Supplemental data, Fig 1). Other researchers have previously found aluminum concentrations to be related to the formation and abundance of sclerotia, primarily in pine forests (Inoue et al., 2006; M. Watanabe et al., 2004; Makiko Watanabe et al., 2004), however none of the soil chemistry data was relatable to the isolate diversity within or between sites. We also observed a much lower abundance of *Cenococcum* sclerotia under *Populus trichocarpa* hosts than reported for other systems. Our methods were only semi-quantitative as we were focused on diversity rather than biomass, however we were typically only able to recover less than 20 sclerotia, and in approximately half our samples zero sclerotia, per gallon (~3.5 kg) of soil sampled. While much of the literature uses similar semi-quantitative methods, one previous study of Douglas Fir forests, also conducted in western Oregon, averaged 0.91 sclerotia per gram of soil (Fogel and Hunt, 1979) translating to 2785 kg ha^−1^ of *Cenococcum* sclerotial biomass. This level of abundance associated with a different host in the same region would seem to be approximately two orders of magnitude greater than those recovered in our study under *Populus*.

### Incongruent genes and indications of recombination within the PNW isolate collection

A side by side visual comparison of the ITS and *GAPDH* phylogenies showed in some cases up to three ITS types associated with a *GAPDH* gene group, and numerous groupings in the respective genes that were inconsistent with each other, a pattern which is strongly suggestive of ongoing or historic intra-species recombination events among the PNW isolates (Fig 3A). Follow on analysis also showed a lack of linear relationships between pairwise distances within the *GAPDH* and ITS datasets (Fig 3B). Further analysis using the inter-partition length difference test (Burt et al., 1996; Swofford, 2003) confirmed that the *GAPDH* and ITS data are incongruent (Fig 3C), indicating that the two genes have conflicting phylogenetic history or else are otherwise drawn from different distributions. These data reflect previous evidence of recombination observed in *C. geophilum* in some of the first studies of this species that used phylogenetic approaches over 20 years ago (LoBuglio and Taylor, 2002) and are largely consistent with studies using a variety of methods both in *Cenococcum* (Bourne et al., 2014; Douhan et al., 2007c, 2007a; Obase et al., 2017) and observed in other ascomycetes (Jurado et al., 2012; Mirete et al., 2013) and interpreted as evidence of cryptic recombination. Similarly, in our study recombination appears to have been active in the PNW isolate collection and is evident in the incongruent histories of inheritance between the ITS and *GAPDH* gene regions. However, while the history of these two genes show patterns consistent with recombination, unfortunately it is just two genes, and gaining evidence across many more loci within the population will be important to solidify this conclusion and fully quantify rate and extent of gene flow.

### Cenococcum phylogeographic diversity within a global context

Our study increases the known diversity of *C. geophilum* within the global isolate collection, with many of the PNW isolate collection clades appearing to be unique to those shown previously (Figure 4). Our analyses also resolved new relationships within the global collection, with isolates tending to group by region or continent of origin, with some notable exceptions where isolates from multiple continents form single well-supported clades. The genetic diversity of our *C. geophilum* isolates from the Pacific Northwest also appears greater in comparison to that found in other regions, with previous studies resolving a maximum of 9 clades within any particular collection (de Freitas Pereira et al., 2018; Douhan et al., 2007a; Douhan and Rizzo, 2005; Matsuda et al., 2015; Obase et al., 2016) (Table 2). Four of the identified clades in the PNW isolates grouped similarly but remained unaffiliated with any of those in the global study, implying that clades three, four, and nine may be specifically associated with *Populus* in the Pacific Northwest or part of a hypothesized “core” of *C. geophilum* isolate collections associated with diverse hosts but just not represented in the sparse global samplings to date. Additionally, our analyses suggest PNW clades 10 and particularly 11 may represent particularly divergent clonal groups or incipient species within the PNW isolate collection, as both clades segregated strongly from the rest of the PNW isolates. This confirms the need for future studies to target representatives from multiple resolved PNW clades for additional genetic analyses using higher resolution genotype-by-sequencing techniques to determine potential speciation within the PNW isolate collection.

Our isolates also add considerably to the known global diversity of this taxon. Four newly designated clades, V16-V19, represent never-before-seen relationships determined through a ML analysis using best fit parameters (Fig 4, Supplemental Data, Table 3). One of these clades, V16, includes isolates from both the North American and European continents, and both clade V8 and V17 include isolates from both the West and East coasts of the United States (Fig 4, Supplemental Data, Table 3). However, much of the topology of the global isolate collection analysis also closely resembles those recovered in previous studies of *C. geophilum* originating across geographically distinct regions. These distinct groups that were present in previous analyses are indicated on the ML analysis using the last initial of the first author (Fig 4, Supplemental data, Table 3). Isolates from Japan grouped in three separate clades, MI, MII and MIII, mirroring the Japanese strain clades I, II and III from Matsuda et al. (2015) (Fig 4, Supplemental data, Table 3). Additionally, four clades were determined in our study which are present, but not strongly supported groupings, in the Matsuda et al. (2015) study: M1, M2, M3, and M4. Each of these consists of 2-3 strongly grouped isolates which also grouped together in the previous study but did not have strong bootstrap support. Clade M1 is supported at bootstrap value 98.8% and includes two French isolates (Fig 4, Supplemental data, Table 3). Clade M2 is supported at bootstrap value 88.3% and includes three Spanish isolates (Fig 4, Supplemental data, Table 3). A second Spanish clade, M3, is supported at bootstrap value 81.8% and includes 3 isolates. Clade M4 includes several subclades of highly supported East Coat, USA isolates (Fig 4, Supplemental data, Table 3).

A 2005 study completed on a collection of *C. geophilum* isolated from Browns Valley, CA in the United States based on *GAPDH*, showed a total of three clades identified at greater that 90% bootstrap support (Douhan and Rizzo, 2005). While isolates from this study collection tended to group together in our analyses as well, the associated clades did not persist save for one nested grouping in clade III, designated here as D2 (Fig 4, Supplemental data, Table 3), which includes 23 isolates at a bootstrap support value of 98.3%.

Obase et al. (2016) previously delineated a total of seven clades in a concatenated *GAPDH* + ITS ML analysis of the Florida/Georgia isolate collection collected with pine trees. The identified Obase et al. clade 7 (O7) was described as the new species designated as *Pseudocenococcum floridanum* gen. et sp. nov. We used this taxon as an outgroup for both the PNW and global studies, as none of our isolates appear to be phylogenetically affiliated with that new species. The remaining relationships described by Obase persist in part in our study and are designated as O1, O2, O3, O4, O5 and O6, but do not directly mirror the previous *GAPDH* ML analysis due to the inclusion of worldwide isolates (Fig 4, Supplemental data, Table 3). Only O3 directly persists from the Obase et al. study to this study as a strongly supported clade at a bootstrap value of 100% (Fig 4, Supplemental data, Table 3), although strong groupings with bootstrap support values >86% exist in O1, O4, O5, and O6 (Supplemental data, Table 3). The isolate sequences provided by Drs. Francis Martin and Martina Peter generally remained unresolved in the global isolate collection analysis, with a few exceptions. Clade dFP1 includes four isolates from Switzerland, including the model genome sequenced strain CG 1.58 (Fig 4, Supplemental data, Table 3). This grouping mirrors the phylogenetic relationship seen between these isolates in de Freitas Pereira et al. (2018). Two other newly designated clades, V18 and V19, include only isolates from Switzerland at bootstrap support values 95% and 98.5%, respectively (Fig 4, Supplemental data, Table 3).

Two of the newly designated clades, V16 and V17, encompass the greatest geographic diversity in the global isolate collection analysis. The clade V16 includes 22 total isolates grouped into 4 strongly supported nested clades (>90%), with three Florida/Georgia isolates, three French isolates, one Swedish isolate, two Polish isolates, and two isolates from Holland comprising the largest nested clade (Fig 4, Supplemental data, Table 3). Clade V16 includes an additional nine Florida/Georgia isolates grouped into two separate nested clades, and two Swiss isolates grouped into a nested clade (Fig 4, Supplemental data, Table 3). Clade V17 (>90%) includes an isolate from Browns Valley, CA, US, and an isolate from the Florida/Georgia isolate collection (Fig 4, Supplemental data, Table 3). PNW clades 3, 4, 9, and 10 show are nearly identical in both RAxML phylogenies, but do not persist in strongly supported clades in the global isolate collection phylogeny (Fig 3, 4). Other isolates which were not strongly grouped in the PNW analysis remained ungrouped in the global isolate collection analysis (Fig 4, Supplemental data, Table 3). Additionally, unresolved isolates persisted in the global isolate collection phylogeny from previous studies (Fig 4, Supplemental data, Table 3).

The large spatial distances, across which several well-resolved clades were observed within both the PNW and global isolate analyses, further highlight the need for higher resolution genetic studies. There is a possibility that greater genetic resolution will subdivide such groups despite spatially distinct origins to strongly group together based on origin. However, the cryptic nature of *C. geophilum* also presents the possibility that these wide distribution patterns between isolates will become more extreme as well. Either case will provide further understanding of the patterns of speciation and genetic exchange within the *C. geophilum* species complex.

### Conclusions and future directions

The genetic diversity present within both local and global isolate collections of *C. geophilum* isolates is striking. Our study reveals the existence of multiple cryptic clades of *C. geophilum* as well as distinct phylogenetic groups from the PNW which appear to date to be uniquely associated with *P. trichocarpa,* and confirms the common view of this species as a hyper-diverse group of ectomycorrhizal fungi on regional and global scales. Our study additionally strongly indicates patterns consistent with recombination within the PNW isolate collection based on analyses performed on the *GAPDH* and ITS gene regions. However, to provide further evidence as to whether this represents one global, hyper-diverse species, or a myriad of cryptic species, a more robust analysis with greater population-level resolution across many loci to more accurately quantify gene flow will be required. Future research could further elucidate these regional and global relationships through the use of genotype-by-sequencing (GBS) or restriction-associated DNA sequencing (RAD-seq) approaches for rigorous *de novo* single nucleotide polymorphism (SNP) analysis (Glenn et al., 2017). Such approaches could help shed a light on this ubiquitous fungal taxon which has proven historically difficult to classify both physiologically and phylogenetically, and allow us the opportunity to delineate the individual species which may currently be included in this greater *C. geophilum* species complex, or the other mechanisms by which it may maintain such diversity.

## Acknowledgements

The authors would like to thank Keisuke Obase, Yosuke Matsuda, Matthew Smith, Francis Martin and Martina Peter for providing access to cultures and sequences for comparison with our isolates, as well as Allison Veach and Stephanie Kivlin for their assistance with statistical analyses. This research was sponsored by the Genomic Science Program, U.S. Department of Energy, Office of Science, Biological and Environmental Research, as part of the Plant Microbe Interfaces Scientific Focus Area at ORNL (http://pmi.ornl.gov). Oak Ridge National Laboratory is managed by UT-Battelle, LLC, for the U.S. Department of Energy under contract DEAC05-00OR22725.

---

**Supplemental Fig 1**: Correlogram of PNW clade, rivershed of origin, site latitude, soil percent moisture content, C/N weight percent, temperature, elemental metal (lead, zinc, copper, cadmium, strontium) counts per minute, total sclerotia isolated, and total *C. geophilum* isolation success of 105 PNW soil samples. Positive correlations are highlighted in blue and negative correlations are highlighted in red, with color intensity proportional to the correlation coefficient. Only those correlation coefficients with p<0.05 are shown in color. No measured soil conditions or qualities were determined to correlate with the total sclerotia obtained or *C. geophilum* successfully isolated.

**Supplemental Table 1**. Pacific Northwest isolate sites and total *Cenococcum geophilum* isolated. The overall isolation success rate of *C. geophilum* from 105 soil samples was 46%. Total *C. geophilum* isolates >0 are bolded in the Total *Cenococcum* column.

**Supplemental Table 2**. Pacific Northwest *Cenococcum geophilum* isolates, rivershed of origin, associated clade and bootstrap support value. Bootstrap values of >85% are bolded. Unresolved isolates or isolates not grouped into a nested clade are designated with a dash (−). The ITS and GAPDH GenBank accession numbers are included.

**Supplemental Table 3**. Global population GAPDH maximum likelihood analysis of 790 *Cenococcum geophilum* strains. Groupings implicated or identified in previous studies are indicate as follows: D (Douhan et al., 2007a; Douhan and Rizzo, 2005); dFP (de Freitas Pereira et al., 2018); M (Matsuda et al., 2015); and O (Obase et al., 2016). Strongly supported clades designated in this study are designated with V. Clades V16-V19 represent newly designated clades within the global *C. geophilum* population. Bootstrap support values of >85% are bolded. Isolates not grouped into a nested clade are designated with a dash (−).

